# Elevated mitochondrial superoxide promotes longevity through a mitochondria-to-nucleus kinase signaling pathway

**DOI:** 10.1101/2025.06.09.658743

**Authors:** Ulrich Anglas, Aura A. Tamez González, Maisha Maliha Promi, Abdelrahman AlOkda, Alain Pacis, Jeremy M. Van Raamsdonk

**Author notes:** **Corresponding Author:** Jeremy Van Raamsdonk, Research Institute of the McGill University Health Centre, 1001 Decarie Blvd., Montreal, Quebec, Canada, 1-514-934-1934 ext. 76157.

## Abstract

The reactive oxygen species superoxide is generated by mitochondria during the process of producing energy. While superoxide can cause oxidative damage to the cell, we and others have shown that a mild increase in mitochondrial superoxide extends longevity in multiple model organisms. To elucidate the molecular mechanisms involved, we identified transcriptional changes in mitochondrial superoxide dismutase deletion mutants (*sod-2* worms) using RNA sequencing. *sod-2* mutants exhibit a number of changes in nuclear gene expression resulting from elevated mitochondrial superoxide suggesting that mitochondria-to-nucleus signaling is contributing to their longevity. Gene ontology enrichment analysis demonstrated that genes involved in innate immunity and cuticle formation are significantly upregulated in *sod-2* worms. To identify kinases involved in this lifespan-extending pathway, we completed a targeted RNA interference screen to examine the contribution of 61 selected kinases to *sod-2* longevity. From this screen, we found 25 kinases which are required for the long lifespan of *sod-2* mutants including *mak-2*, which has an established role in a kinase signaling pathway involved in axon regeneration. Disruption of *mak-2* specifically reduces *sod-2* lifespan but not wild-type longevity and also decreases resistance to multiple exogenous stressors. In examining other genes that act with *mak-2* in pathways controlling axon regeneration, we identified a SEK-3/PMK-3/MAK-2/CEBP-1 signaling pathway that is specifically required for *sod-2* longevity but not wild-type lifespan. Combined these results suggest a novel role for kinases with established roles in axon regeneration in promoting longevity through a mitochondria-to-nucleus signaling pathway.

## Introduction

Superoxide is the main form of reactive oxygen species (ROS) that is generated in the mitochondria when electrons from the electron transport chain react directly with oxygen. As ROS can cause oxidative damage to multiple components of the cell, including DNA, protein and lipids, the Free Radical Theory of Aging (FRTA) proposes that aging is caused by the accumulation of oxidative damage resulting from ROS generated by normal metabolism (1). The mitochondrial FRTA further proposes that the accumulation of oxidative damage in the mitochondria decreases the efficiency of the electron transport leading to an accelerated generation of ROS in a self-amplifying cycle.

In order to control the levels of ROS, organisms express antioxidant enzymes, which act to detoxify ROS. Superoxide dismutase (SOD) converts superoxide to oxygen and hydrogen peroxide. Humans have three different SOD proteins, named SOD1, SOD2 and SOD3, which are located in the cytosol, mitochondria and extracellular space, respectively. In addition to the cytoplasmic (SOD-1), mitochondrial (SOD-2) and extracellular (SOD-4) SOD proteins, *C. elegans* has two additional SOD proteins, SOD-3 and SOD-5, which are normally expressed at low levels in the mitochondria and cytoplasm, respectively.

We and others have previously examined the effect of disrupting *sod* genes on the lifespan of *C. elegans.* While the FRTA would predict that reducing antioxidant defense would result in decreased lifespan, individually disrupting the five *C. elegans sod* genes had little or no detrimental effect on longevity, despite increasing sensitivity to oxidative stress (2–4). In fact, deletion of all five *sod* genes in the same worm did not decrease lifespan compared to wild-type animals, indicating that superoxide dismutase is not required for normal lifespan under unstressed conditions (5).

Interestingly, deletion of the mitochondrial *sod* gene *sod-2* resulted in increased lifespan (3). This increase in lifespan is believed to be caused by elevated mitochondrial superoxide levels resulting from decreased superoxide detoxification. This conclusion is supported by the fact that directly increasing superoxide levels through treatment with small concentrations of the superoxide generating compounds paraquat or juglone can also extend longevity, while reducing superoxide levels in *sod-2* worms with antioxidants decreases their lifespan (6, 7).

At present, the mechanisms by which elevated mitochondrial superoxide levels lead to increased lifespan are incompletely elucidated. We hypothesized that increased levels of mitochondrial superoxide act as a stress signal that is communicated to the nucleus to change genes expression, and that while these changes in gene expression are activated to survive the superoxide stress they also lead to lifespan extension. To test this hypothesis, we used RNA sequencing (RNA-seq) to examine gene expression in *sod-2* mutants compared to wild-type and show that mitochondrial perturbations in *sod-2* worms result in widespread changes in nuclear gene expression. To elucidate the mechanism by which mitochondrial stress is communicated to the nucleus, we completed a targeted RNA screen that identified a number of kinases that are required for *sod-2* longevity. Among the kinases identified was MAK-2, a kinase that is part of an established signaling pathway that promotes axon regeneration. We further find that other proteins involved in this axon regeneration signaling pathway are also required for *sod-2* lifespan suggesting that a SEK-3/PMK-3/MAK-2/CEBP-1 signaling cascade is involved in the mitochondria-to-nucleus signaling pathway that promotes longevity in *sod-2* worms.

## Results

### Disruption of mitochondrial superoxide dismutase results in changes in nuclear gene expression

To gain insight into the molecular mechanisms contributing to lifespan extension in *sod-2* mutants, we performed RNA-seq on day 1 and day 8 adult *sod-2* and wild-type worms. A principal component analysis (PCA) demonstrated that there is a clear separation in gene expression between *sod-2* and wild-type worms at day 1 of adulthood, with very little difference at day 8 of adulthood (**Figure 1A**). This conclusion is supported by visualization of gene expression using heat maps (**Figure 1B,C**). Interestingly, we observed much larger changes in gene expression between day 1 and day 8 animals than between the two genotypes (**Figure S1**). At day 1 of adulthood, we found 775 genes that were significantly upregulated in *sod-2* worms compared to wild-type and 125 genes that were significantly downregulated (**Figure 1D; Table S1**). In contrast, at day 8 of adulthood, we only observed 17 and 12 genes that were significantly upregulated or downregulated, respectively, in *sod-2* worms compared to wild-type (**Figure 1D; Table S1**). Comparing between the time points, there were 10 upregulated genes and 5 downregulated genes in common (**Figure 1E,F**; **Table S1**). Overall, these findings show that disruption of *sod-2* in the mitochondria causes widespread changes in nuclear gene expression.

**Figure 1.**
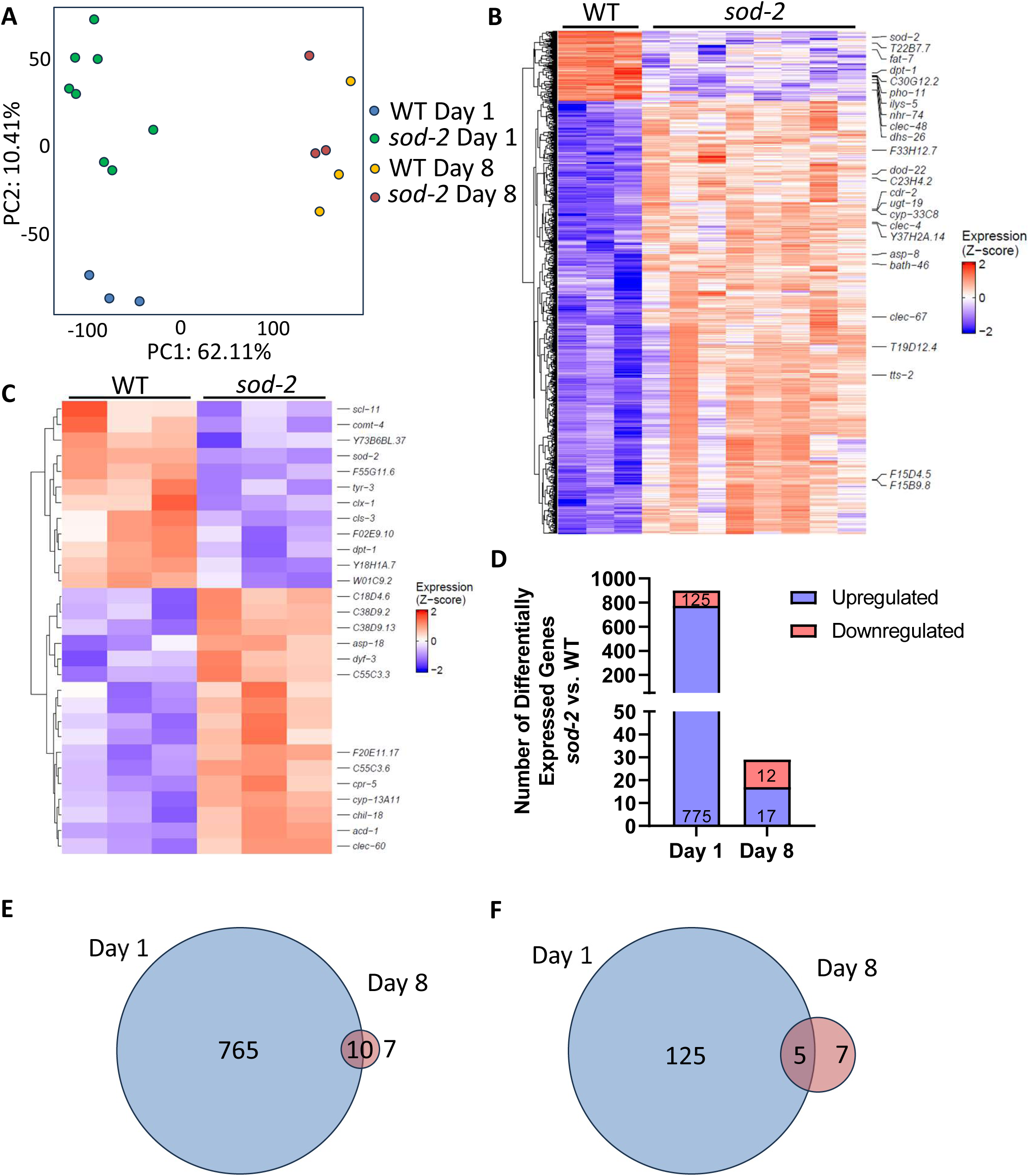
Elevated mitochondrial superoxide results in changes in nuclear gene expression. RNA sequencing was used to compare gene expression between wild-type and *sod-2* worms on day 1 and day 8 of adulthood. **(A)** The principal component analysis (PCA) plot shows a clear separation between the two genotypes at day 1 of adulthood and between day 1 and day 8 samples. **(B)** A heat map comparing gene expression between *sod-2* and wild-type worms at day 1 of adulthood shows a number of significantly upregulated genes in *sod-2* worms and a lesser number of significantly downregulated genes. **(C)** A heat map showing differentially expressed genes between wild-type and *sod-2* worms at day 8 of adulthood. **(D)** At day 8 of adulthood, there were much fewer differentially expressed genes in *sod-2* worms compared to wild-type worms than at day 1 of adulthood. Comparing differentially expressed genes at the two times points shows that 10 upregulated genes **(E)** and 5 downregulated genes **(F)** are significantly differentially expressed in *sod-2* worms at both time points.

Having identified differentially expressed genes in *sod-2* mutants, we next performed enrichment analyses to identify over-represented pathways. Using the ***G****ene* ***O****ntology en****RI****chment ana****L****ysis and visua****L****iz****A****tion tool* (GOrilla) and ShinyGO to perform the enrichment analysis, we found that the main biological processes that were overrepresented among the genes upregulated in *sod-2* worms at day 1 of adulthood were cuticle development and innate immune stress response (**Figure S2, S3**). Both of these processes have been previously associated with longevity and would serve to protect worms against pathogens (8–10). The increase in ROS resulting from the loss of *sod-2* may mimic exposure to bacterial pathogens, which also increases ROS (11) and leads to the activation of innate immune signaling pathways.

### Targeted RNAi screen identifies kinases required for long lifespan of *sod-2* mutants

Having shown that mitochondrial ROS alters nuclear gene expression in *sod-2* mutants, we hypothesized that kinase signaling is involved in transmitting a signal from mitochondria to the nucleus to indicate that mitochondrial superoxide levels are elevated (**Figure 2A**). To identify kinases involved in this mitochondria-to-nucleus signaling pathway, we performed a targeted RNA interference (RNAi) screen (**Figure 2B**). We used RNAi to knock down kinases in *sod-2* worms beginning at the L4 stage of the parental generation and measured the lifespan of the progeny. In total, we measured the lifespan of *sod-2* worms treated with RNAi targeting 61 different, prioritized kinases, with at least three biological replicates per RNAi clone, and identified 25 kinases that significantly decrease *sod-2* lifespan when they are disrupted (**Figure 2C; Figure S4**).

**Figure 2.**
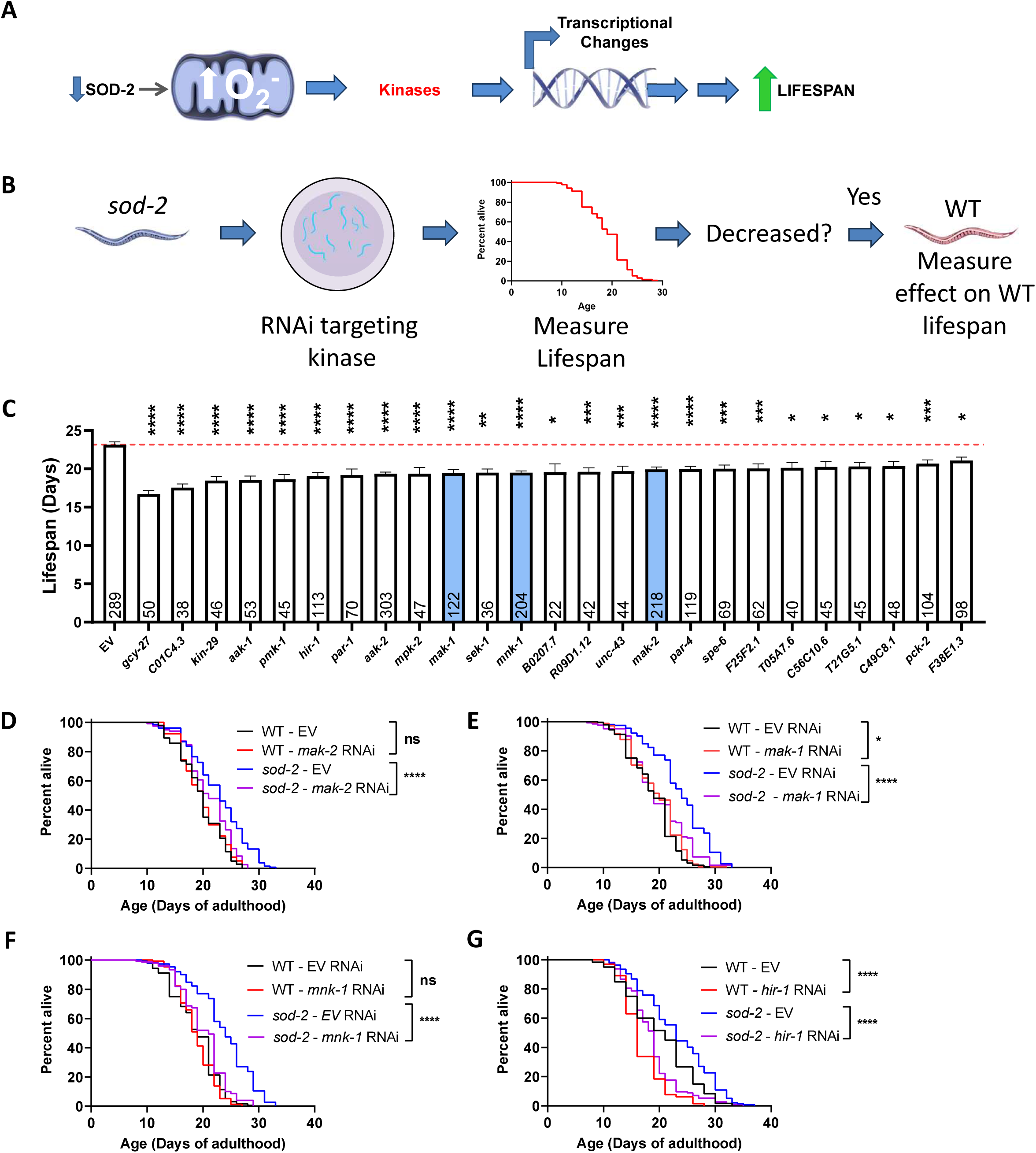
Targeted RNA interference screen identifies multiple kinases contributing to lifespan extension. **(A)** Working model for how *sod-2* deletion extends longevity: increased mitochondrial superoxide leads to activation of cytoplasmic kinase signaling pathways, which activate transcription factors to change gene expression and increase lifespan. **(B)** To identify kinases required for *sod-2* longevity, we completed a targeted RNAi screen. **(C)** The screen identified multiple kinases that are required for *sod-2* longevity. Of 61 kinase RNAi clones tested, 25 RNAi clones significantly decreased *sod-2* lifespan. All lifespans in panel C were performed in *sod-2* worms. Red line indicates the lifespan of *sod-2* worms on empty vector (EV) bacteria. Disruption of *mak-2* **(D)***, mak-1* **(E)** or *mnk-1* **(F)** significantly decrease *sod-2* lifespan but do not decrease wild-type longevity. **(G)** In contrast, *hir-1* RNAi was generally toxic decreasing both wild-type and *sod-2* lifespan. Error bars indicate SEM. Three biological replicates were completed. Statistical analysis was performed using a one-way ANOVA with Dunnett’s multiple comparisons test in panel C and a log-rank test in panels D-G. ns = not significant, *p<0.05, **p<0.01, ***p<0.001, ****p<0.0001.

Among the kinases found to be required for *sod-2* lifespan, were kinases that have been previously implicated in longevity. For example, knockdown of *sek-1* or *pmk-1* from the p38-mediated innate immune signaling pathway reduced *sod-2* lifespan and have previously been shown to be required for the extended longevity of other long-lived mutants (9, 10, 12–14). Similarly, both *aak-1* and *aak-2*, the genes encoding AMP-activated protein kinase (AMPK), and *par-4*, which encodes the *C. elegans* homolog of the LKB1 kinase that activates AMPK, are required for *sod-2* lifespan, and have previously been shown to be required for the long lifespan of other long-lived mutants (15–17). The identification of kinases previously associated with longevity provides validation for the RNAi screen.

Among the kinases found to be required for *sod-2* longevity, was MAK-2 (MAP kinase activated protein kinase 2), a kinase with an established role in a signaling pathway promoting axonal regeneration, and its two paralogs MAK-1 and MNK-1. As little is known about the role of these kinases in longevity, we decided to focus the remainder of our study on MAK-2. Before proceeding with a more in-depth characterization of the role of MAK-2 in the phenotype of *sod-2* mutants, we first wanted to determine whether the effect of *mak-2, mak-1* and *mnk-1* knockdown on lifespan is specific to *sod-2* mutants or whether RNAi targeting these genes also decreases wild-type lifespan. To do this, we knocked down each of the three kinases in both wild-type and *sod-2* backgrounds and found that knocking down any of these kinases reduces *sod-2* lifespan but does not impact wild-type lifespan (**Figure 2D-F**). This indicates that *mak-2, mak-1* or *mnk-1* are specifically required for the extended longevity of *sod-2* worms but are dispensable for normal lifespan in wild-type worms. In contrast, we found that disruption of other kinases, such as *hir-1*, decreased both wild-type and *sod-2* lifespan suggesting a more general contribution to longevity (**Figure 2G**).

### *mak-2* is required for mitochondrial superoxide to increase lifespan

To further confirm the results of the RNAi screen, we examined the effect of a *mak-2* deletion on *sod-2* lifespan. We crossed *sod-2* worms to the *mak-2(gk1110)* deletion mutant, which has a 1024 base pair deletion in the *mak-2* gene combined with the insertion of one thymine. Similar to our results using RNAi, we found that deletion of *mak-2* specifically decreased the lifespan of *sod-2* mutants but had no effect on the longevity of wild-type worms (**Figure 3A**). The lifespan of *sod-2;mak-2* double mutants was equivalent to wild-type. This confirms that results of the RNAi lifespan experiment and indicates that the effect of *mak-2* RNAi on *sod-2* lifespan is due to the knocking down *mak-2* expression, not potential off-target effects.

**Figure 3.**
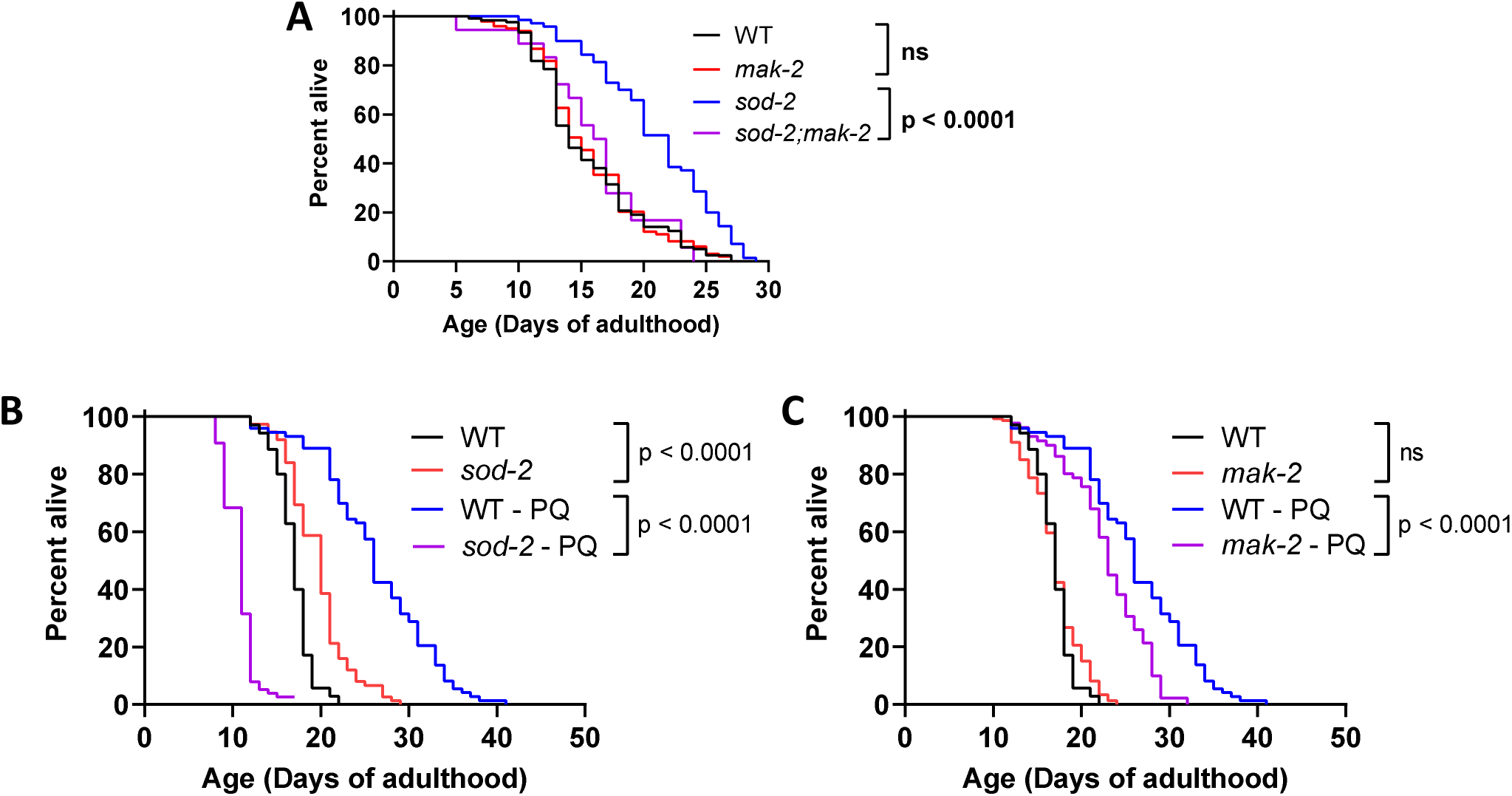
MAK-2 kinase is required for lifespan extension resulting from elevated mitochondrial superoxide. **(A)** Deletion of *mak-2* decreases *sod-2* lifespan but has no effect on wild-type longevity. **(B)** Worms were treated with 0.1 mM paraquat to increase mitochondrial superoxide levels. This concentration of paraquat greatly increases wild-type lifespan, while markedly decreasing the lifespan of long-lived *sod-2* mutants. Disruption of *mak-2* **(C)** decreases the ability of 0.1 mM paraquat treatment to increase lifespan. Three biological replicates were performed. Statistical significance was assessed using a log-rank test. ns = not significant.

Similar to deletion of *sod-2*, directly increasing mitochondrial superoxide levels through treatment with low concentrations of the superoxide generating compound paraquat increases lifespan (6, 18). To determine whether *mak-2* is also required for low doses of paraquat to extend lifespan, we treated wild-type and *mak-2* worms with 0.1 mM paraquat, a concentration that we previously found to be optimal for promoting longevity (18). As a control, we also treated *sod-2* worms with 0.1 mM paraquat, as we have previously shown that even low concentrations of paraquat are detrimental for *sod-2* worms.

We found that 0.1 mM paraquat markedly decreased the lifespan of *sod-2* worms (**Figure 3B**) indicating that the paraquat treatment was effectively increasing superoxide levels. As we have previously observed, 0.1 mM paraquat significantly increased the lifespan of wild-type worms (**Figure 3B**). When *mak-2* deletion mutants were treated with 0.1 mM paraquat, the magnitude of lifespan extension was significantly decreased compared to 0.1 mM paraquat-treated wild-type worms (**Figure 3C**). Combined, this indicates that *mak-2* is required for lifespan extension resulting from the mild increase of mitochondrial superoxide levels.

### Disruption of *mak-2* affects resilience and physiologic rates

Having shown that *mak-2* is required for the long lifespan of *sod-2* mutants, we next sought to determine whether *mak-2* affects other phenotypes in *sod-2* worms including stress resistance and physiologic rates. We examined the effect of *mak-2* deletion on resistance to six different exogenous stressors in wild-type and *sod-2* worms including heat stress (37°C), osmotic stress (450 mM NaCl), acute oxidative stress (320 µM juglone), chronic oxidative stress (4 mM paraquat), anoxia (48 hours) and bacterial pathogen stress (*P. aeruginosa* strain PA14).

As we have previously observed (13), *sod-2* worms have increased heat stress resistance compared to wild-type worms and this enhanced survival was lost when *mak-2* is disrupted (**Figure 4A**). In contrast, deletion of *mak-2* did not significantly affect heat stress resistance in wild-type worms. In the osmotic stress assay, deletion of *mak-2* significantly decreased the survival of both wild-type and *sod-2* worms (**Figure 4B**). This indicates that MAK-2 is generally required for surviving osmotic stress.

**Figure 4.**
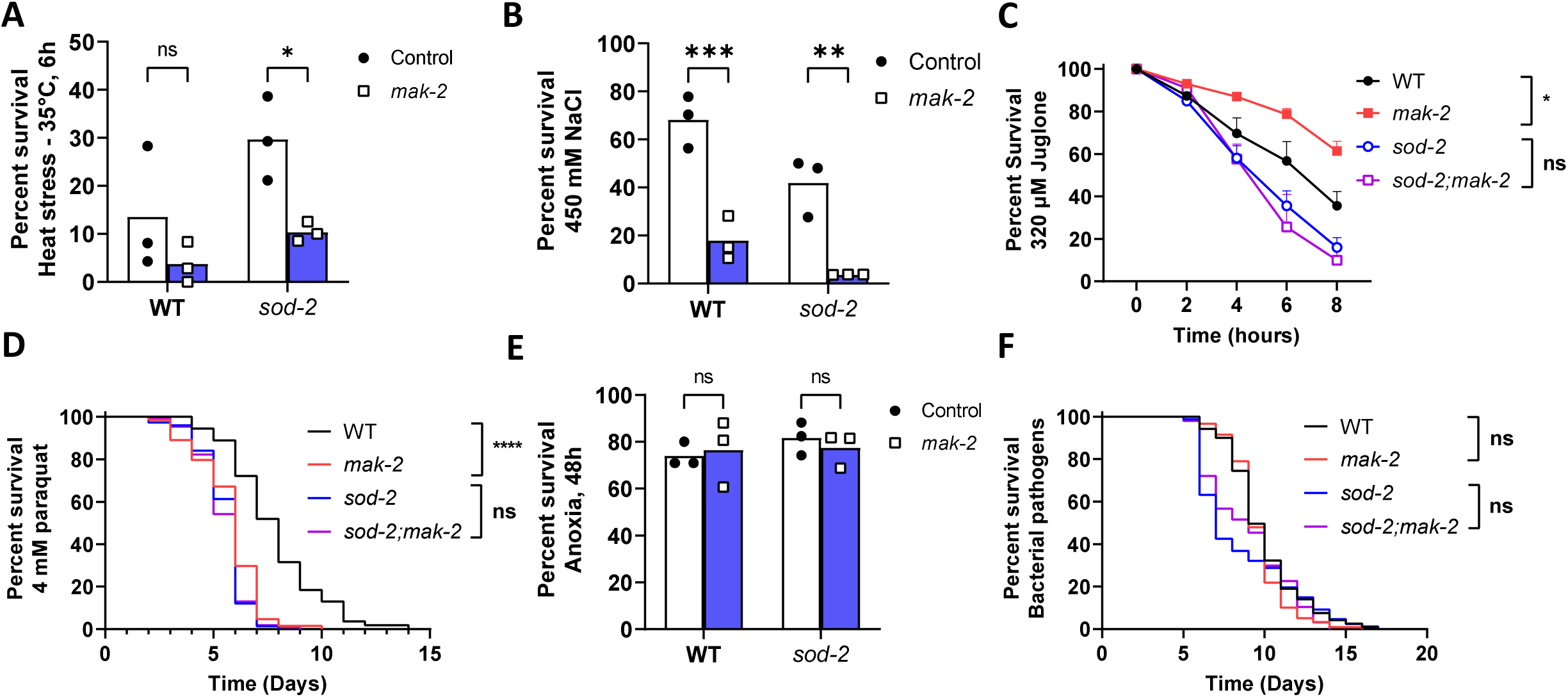
The *mak-2* kinase affects resistance to stress in wild-type and *sod-2* mutants. Loss of *mak-2* decreases resistance to heat stress (**A**; 35°C, 6 hours) and osmotic stress (**B**; 450 mM NaCl) in *sod-2* worms. Deletion of *mak-2* increases resistance to acute oxidative stress (**E**; 320 µM juglone) in wild-type worms but has no effect in *sod-2* mutants. *mak-2* deletion mutants have decreased resistance to chronic oxidative stress (**D**; 4 mM paraquat) but loss of *mak-2* does not further decrease paraquat resistance in *sod-2* worms. Disruption of *mak-2* does not affect resistance to anoxia (**E**; 48 hours) or bacterial pathogens (**F**; *P. aeruginosa* strain PA14, slow kill assay). Error bars indicate SEM. Three biological replicates were completed. Statistical analysis was performed using a log-rank test in panels D and F, a two-way ANOVA with Šidák’s multiple comparisons test in panels A, B and E and a two-way repeated measures ANOVA with Bonferroni’s multiple comparisons test in panel C. ns = not significant, *p<0.05, **p<0.01, ***p<0.001, ****p<0.0001.

Our previous work has shown that *sod-2* worms have decreased resistance to both acute and chronic oxidative stress (3, 13), presumably resulting from the diminished ability to detoxify superoxide. In examining the role of *mak-2* in *sod-2* worms’ resistance to oxidative stress, we found that deletion of *mak-2* had no effect on either acute or chronic oxidative stress survival in *sod-2* worms (**Figure 4C,D**). Interestingly, *mak-2* mutant worms had significantly higher levels of survival than wild-type worms in the acute oxidative stress assay, but significantly decreased survival in the chronic oxidative stress assay.

Finally, we examined the role of *mak-2* in surviving anoxia and bacterial pathogen stress. In both cases, we found that disruption of *mak-2* did not affect the survival of either wild-type or *sod-2* worms (**Figure 4E,F**). Overall, these results indicate that MAK-2 is required for the enhanced heat stress resistance of *sod-2* worms, contributes to chronic oxidative stress resistance in wild-type worms and is generally required for osmotic stress resistance in wild-type and *sod-2* worms.

We next examined the effect of *mak-2* deletion on physiologic rates, including fertility, development and movement, all of which can be viewed as measures of general health. The brood size of *sod-2* worms is decreased compared to wild-type animals but not altered by the disruption of *mak-2* (**Figure 5A**). *mak-2* mutants exhibit a small reduction in brood size compared to wild-type worms (**Figure 5A**). The slow post-embryonic development time of *sod-2* worms is further slowed by the loss of *mak-2*, which does not lengthen the development time of wild-type worms (**Figure 5B**). Disruption of *mak-2* resulted in a small decrease in the thrashing rate of both wild-type and *sod-2* worms. Thus, *mak-2* deletion specifically slows down development in *sod-2* worms, but inhibits movement in both strains.

**Figure 5.**
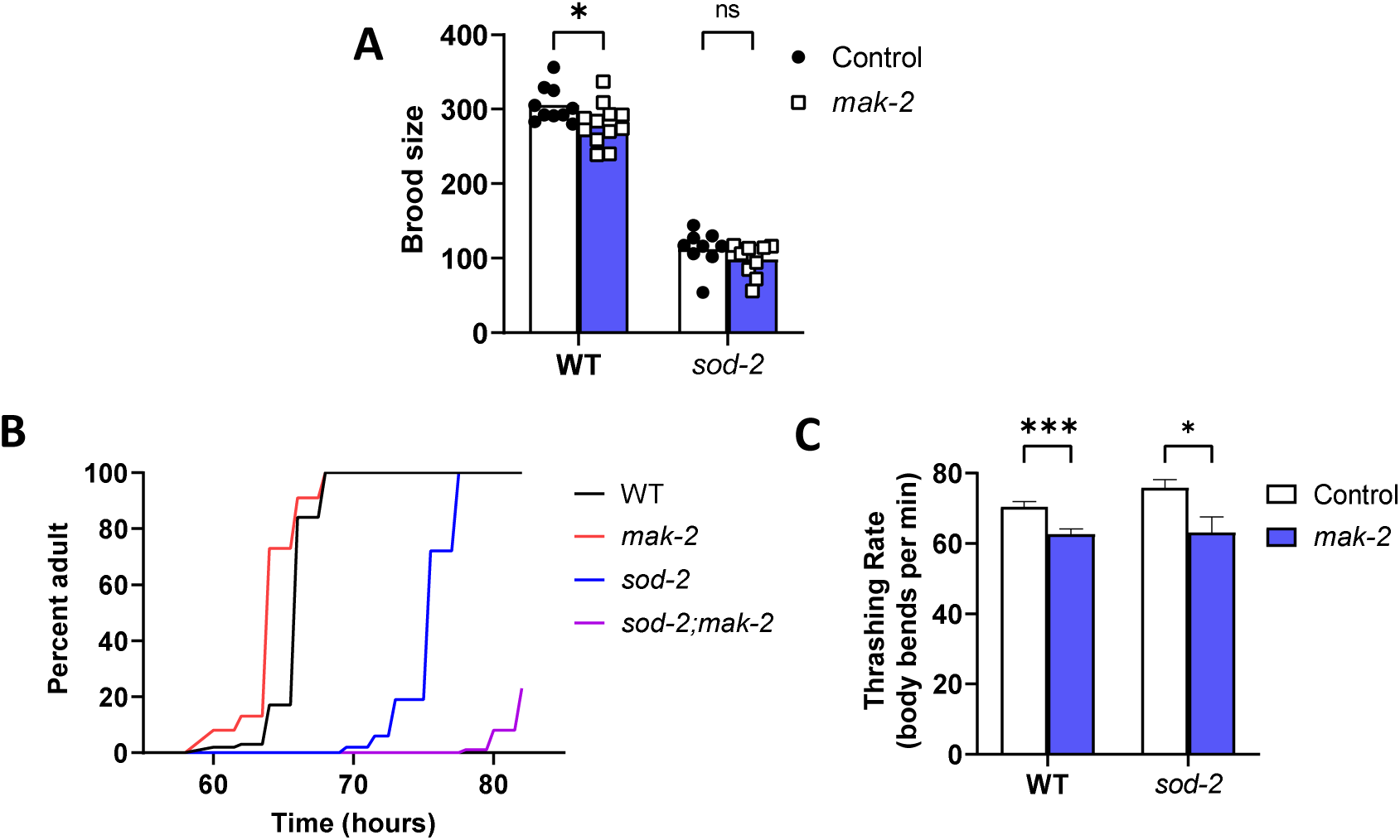
Disruption of *mak-2* kinase exacerbates the slow development of *sod-2* mutants. **(A)** Disruption of *mak-2* slightly reduces fertility in wild-type worms but does not further decrease brood size in *sod-2* mutants. **(B)** *mak-2* deletion slows post-embryonic development time in *sod-2* worms but does not delay development in wild-type animals. **(C)** Loss of *mak-2* results in a small decrease in thrashing rate in both wild-type and *sod-2* worms. Error bars indicate SEM. Three biological replicates were completed. Statistical analysis was performed using a two-way ANOVA with Šidák’s multiple comparisons test in panels A and C, and a log-rank test in panel B,. ns = not significant, *p<0.05, ***p<0.001.

### Role of axonal regeneration pathways in *sod-2* longevity

While MAK-2 has not previously been associated with longevity, the MAK-2 kinase has an established role in the DLK-1 kinase signaling pathway that promotes axon regeneration (19–22). In this pathway, DLK-1 activates MKK-4, which activates PMK-3. PMK-3 then activates MAK-2, which activates the transcription factor CEBP-1. In parallel to the DLK-1 pathway, the MLK-1 pathway involving the kinases MLK-1, MEK-1, and KGB-1 as well as the transcription factor FOS-1 has also been shown to contribute to axon regeneration (19–22). There is also a great deal of cross-communication between these pathways. The phosphatases VHP-1, PPM-1 and PPM-2 as well as the E3 ubiquitin ligase RPM-1 can all inhibit both the DLK-1 and MLK-1 pathways (20, 21, 23–26). To determine whether axon regeneration pathway genes other than *mak-2* are required for the long lifespan of *sod-2* mutants, we used RNAi to knock down the expression of these genes in both *sod-2* and wild-type worms and compared the lifespan to worms grown on empty vector (EV) control bacteria.

While the knockdown of *dlk-1* or *mkk-4* did not decrease *sod-2* longevity, RNAi targeting *pmk-3, mak-2* or *cebp-1* all specifically reduced *sod-2* lifespan without affecting wild-type lifespan (**Figure 6A-D**, **Figure 2D**). As SEK-3 has been proposed to act in a pathway with PMK-3 (27), we also examined the effect of *sek-3* RNAi and found that it specifically reduced the lifespan of *sod-2* mutants but not wild-type worms (**Figure S5**). Treatment with RNAi targeting *rpm-1* also specifically decreased the lifespan of *sod-2* worms but not wild-type worms (**Figure 2E**). In the MLK-1 pathway, decreasing the expression of either *mlk-1* or *mek-1* did not decrease the lifespan of either *sod-2* or wild-type worms, though interestingly *mlk-1* RNAi significantly increased lifespan (**Figure 6E,F**). For several of the genes involved in the axon regeneration pathways, including *kgb-1, fos-1, vhp-1, ppm-1* and *ppm-2,* we found that disruption of the genes with RNAi decreased both wild-type and *sod-2* lifespan, indicating that these genes are generally required for lifespan (**Figure 6H-L**). Overall, these results suggest the possibility that a kinase signaling pathway involving SEK-3/PMK-3/MAK-2/CEBP-1 is specifically required for lifespan extension in long-lived *sod-2* worms but not needed for wild-type longevity (**Figure 6M**).

**Figure 6.**
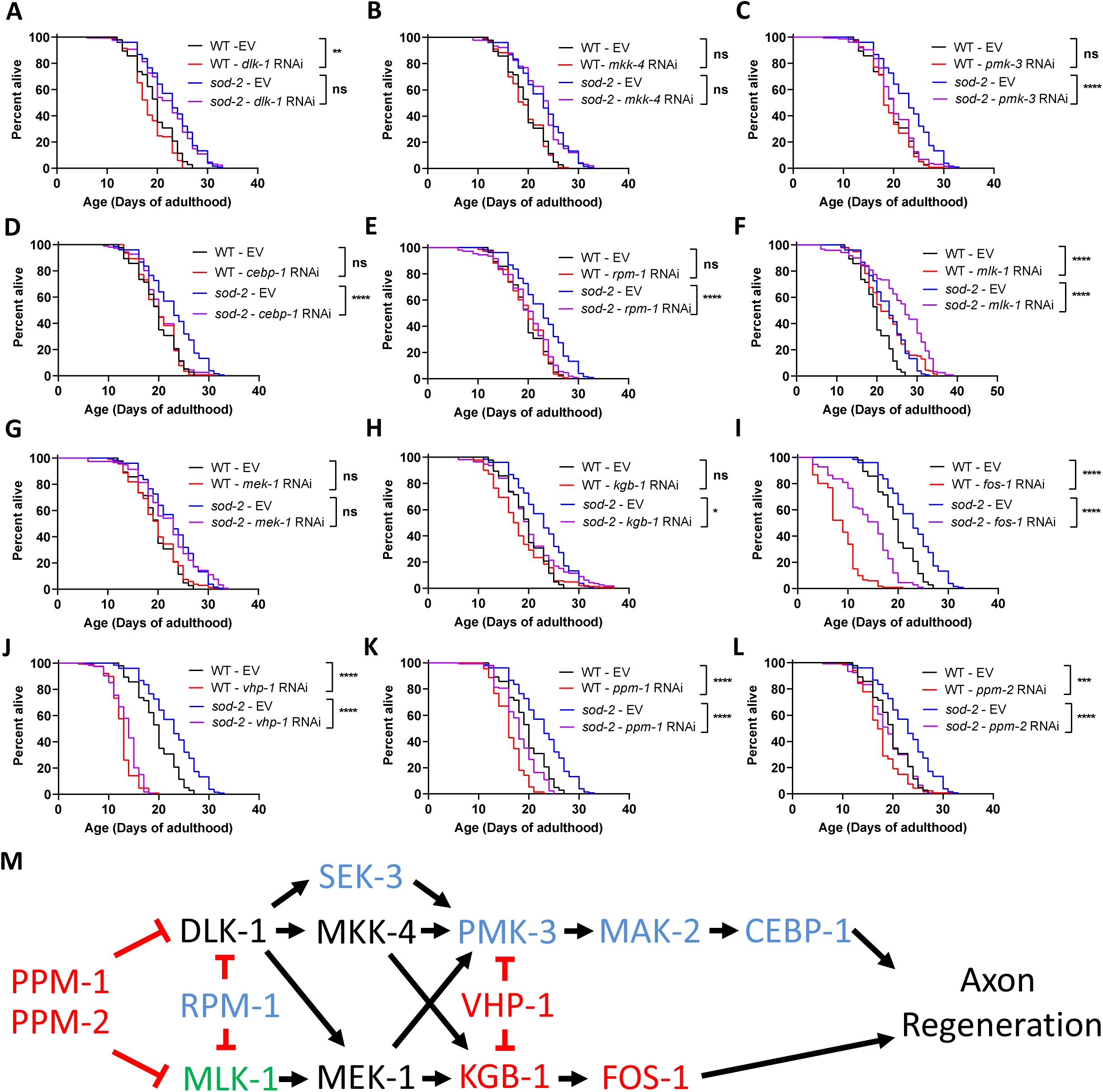
A SEK-3/PMK-3/MAK-2/CEBP-1 signaling pathway is specifically required for lifespan extension in *sod-2* mutants. Based on MAK-2’s established role in pathways contributing to axon regeneration, we evaluated the role of these pathways in the longevity of *sod-2* mutants. RNA interference was used to decrease the expression of genes involved in axon regeneration pathways. *dlk-1* **(A)***, mkk-4* **(B)**, and *mek-1* **(F)** were not required for longevity. Disruption of *pmk-3* **(C)***, mak-2* (Figure 2D)*, cebp-1* **(D)** or *rpm-1* **(I)** decreased *sod-2* lifespan but did not affect wild-type lifespan. This suggests that these genes specifically contribute to the extended longevity of *sod-2* worms. Disruption of *mlk-1* increased both wild-type and *sod-2* lifespan **(E)**. Knockdown of *kgb-1* **(G)***, fos-1* **(H)***, vhp-1* **(J)***, ppm-1* **(K)** or *ppm-2* **(L)** decreased both wild-type and *sod-2* lifespan suggesting that these genes are generally required for lifespan. **(M)** Summary diagram of effects of axon regeneration pathway genes on longevity. Black text = no effect on lifespan. Blue text = specifically required for extended longevity of *sod-2* worms. Red text = disruption decreases wild-type and *sod-2* lifespan. Green text = knockdown results in increased lifespan. Three biological replicates were performed. Statistical significance was assessed using a log-rank test.

To determine the extent to which CEBP-1 target genes are upregulated in *sod-2* mutants, we compared genes upregulated in *sod-2* worms on day 1 of adulthood to genes bound by CEBP-1. The list of genes bound by CEBP-1 was previously identified using Chromatin Immunoprecipitation Sequencing (ChIP-seq) in worms expressing FLAG-tagged CEBP-1 in a *cebp-1* mutant background under control conditions (28). We identified 16 genes in the overlap between the two gene sets (**Figure S6A**). This represents a 1.9 fold enrichment compared to the overlap predicted if two similarly-sized gene sets were chosen at random and is statistically significant (p<0.009). The level of expression of these CEBP-1 target genes ranges from 255% to 6824% in *sod-2* mutants (**Figure S6B**). Among the CEBP-1 target genes that are upregulated are *sqst-1*, which is involved in autophagy, *tbb-6,* which is involved in the unfolded protein response, and multiple genes involved in innate immunity (F56D2.5, K09D9.1, T19D12.4). Interestingly, the expression of *cebp-1* exhibits a trend towards increase in *sod-2* worms (**Figure S6C**).

Finally, we examined the extent to which the upregulation of the CEBP-1 target genes in *sod-2* worms is dependent on the SEK-3/PMK-3/MAK-2 signaling pathway. We found that disruption of *sek-3, pmk-3* or *mak-2* using RNAi decreased the expression of multiple CEBP-1 target genes in *sod-2* worms (**Figure S7**). In cases where we did not observe a decrease in expression after RNAi treatment, it is unclear whether the RNAi knockdown was not sufficient to reduce gene expression or whether other pathways are driving the upregulation of those CEBP-1 target genes. Overall, these results suggest that a kinase signaling pathway involving SEK-3, PMK-3, MAK-2 and CEBP-1 contributes to the longevity of *sod-2* worms at least partially through the upregulation of genes involved in innate immunity and autophagy.

## Discussion

In this paper, we explore the mechanisms underlying lifespan extension in long-lived *sod-2* worms. RNA-seq demonstrates that disruption of superoxide dismutase in mitochondria results in widespread changes in gene expression including upregulation of genes involved in cuticle development and innate immunity. To understand how the disruption of mitochondrial superoxide dismutase is communicated to the nucleus we completed a targeted RNAi screen to identify a MAPK kinase signaling pathway that is specifically required for the long lifespan of *sod-2* mutants but not needed for the longevity of wild-type worms.

### Transcriptional changes in long-lived *sod-2* mutants

To gain insight into the molecular mechanisms underlying lifespan extension in *sod-2* mutants, we performed RNA-seq. Our RNA-seq analysis identified a number of transcriptional changes between *sod-2* and wild-type worms at day 1 of adulthood (900 differentially expressed genes) and far fewer changes at day 8 of adulthood (29 differentially expressed genes). Among the significantly upregulated genes there was an enrichment of genes involved in cuticle development and innate immunity. Previous work has demonstrated a role for the cuticle in longevity (8). The cuticle provides a barrier between the worm and its environment. It protects the worm from physical damage, provides a barrier to inhibit pathogens, prevents internal molecules from exiting the worm and inhibits external compounds from entering the worm (29).

Enhancement of innate immunity would serve to protect the worms against various pathogens. A number of long-lived *C. elegans* mutants show increased expression of genes involved in innate immunity and increased resistance to bacterial pathogens (13). Interestingly, there is a highly significant degree of overlap between genes correlated with lifespan and genes correlated with bacterial pathogen resistance, suggesting that many of the same genetic pathways contribute to both phenotypes (13). The importance of innate immunity for *sod-2* lifespan is demonstrated by the fact that the p38-mediated innate immune signaling pathway is required for their long lifespan, as deletion of *pmk-1* or *sek-1* RNAi both decrease *sod-2* lifespan (13). Similarly, disruption of genes in the p38-mediated innate immune signaling pathway have been shown to decrease the lifespan of multiple other long-lived mutants (9, 10, 12, 13, 30), indicating an important role of innate immunity in determining longevity.

### Kinases contributing to longevity in *sod-2* mutants

In our targeted RNAi screen to identify kinases required for *sod-2* lifespan, we identified 25 kinases that significantly decrease *sod-2* lifespan when they are disrupted. While the majority of these genes have not been previously associated with longevity, our results identified kinases from two pathways known to contribute to longevity: the p38-mediated innate immune signaling pathway (*sek-1, pmk-1*) (9, 10, 12–14) and AMPK (*aak-1, aak-2, par-4*) (15–17), thereby validating our approach to identifying kinases contributing to longevity. Consistent with our results showing the importance of AMPK for *sod-2* lifespan, the AMPK pathway was previously shown to be required for the lifespan extension resulting from treatment with low concentrations of paraquat (17).

Because these other pathways had been well studied with respect to lifespan, we chose to focus our subsequent experiments on MAK-2 and its novel role in longevity. For most of the genes in the RNAi screen, we did not test the effect of knocking down the kinase on lifespan in wild-type worms. Thus, it is unclear whether the other kinases that were required for *sod-2* longevity are generally required for lifespan, as we observed with HIR-1, or specifically required for *sod-2* lifespan, as we observed for MAK-2, MAK-1 and MNK-1. Nonetheless, our screen highlights the importance of kinase signaling in lifespan determination.

Prior to our study, one of the main established roles of MAK-2 was in axon regeneration. MAK-2 acts in a DLK-1 signaling pathway involving MKK-4, PMK-3, MAK-2 and CEBP-1. Interestingly, we found that *pmk-3, mak-2* and *cebp-1* are specifically required for *sod-2* lifespan while disrupting upstream members of the DLK-1 pathway (*dlk-1, mkk-4*) does not affect *sod-2* longevity. It is possible that the RNAi clones targeting *dlk-1* and *mkk-4* did not knock down the levels of *dlk-1* and *mkk-4* sufficiently to affect lifespan. Alternatively, it is possible that other upstream kinases are involved in the activation of the PMK-3/MAK-2/CEBP-1 pathway involved in longevity. As previous work implied the possibility that SEK-3 can activate PMK-3 (27), we tested *sek-3* RNAi and found that *sek-3* is also specifically required for *sod-2* longevity. Combined, these results suggest the possibility that a SEK-3/PMK-3/MAK-2/CEBP-1 signaling pathway is contributing to longevity.

At least some members of this pathway have been previously associated with longevity. Mutation of *sek-3* decreases the extended longevity resulting from knockdown of *isp-1* with RNAi (27). The loss of *pmk-3* was previously shown to be required for lifespan extension resulting from *atp-3* RNAi, *isp-1* point mutation, *isp-1* RNAi, *tpk-1* mutation, *nuo-2* RNAi, and *cco-1* RNAi, but expendable for the long lifespan of *nuo-6* and *clk-1* mutants (27). The loss of *pmk-3* has also been found to further decrease the shortened lifespan of LonP1 protease (*lonp-1*) mutants (31). Consistent with a role for PMK-3 signaling in longevity, disruption of the VHL-1 phosphatase gene *vhl-1* results in activation of PMK-3 and increased lifespan (32), though the increase in lifespan was not directly shown to result from PMK-3 activation. Finally, it has been shown that *mak-2* RNAi prevents the lifespan extension resulting from *atp-3* RNAi when *atm-1* or *atl-1* are also disrupted, but not in wild-type worms (33).

In addition to axon regeneration, components of the DLK-1 signaling pathway have been shown to have other roles in the organism such as controlling cilia length (34), inducing autophagy in response to neuronal injury (35), decreasing protein expression in response to microtubule disruption (36), activating the ER stress response (37), inhibiting axon degeneration in response to loss of mitochondria (38), and so forth. It is unclear whether the role of DLK-1 pathway signaling components in axon regeneration or these other functions contributes to its impact on the longevity of *sod-2* mutants, or whether this represents a completely novel function of this pathway.

### Mechanisms of lifespan extension in *sod-2* mutants

Our results demonstrate that *sod-2* longevity is dependent on the CEBP-1 transcription factor and that targets of the CEBP-1 transcription factor are enriched among the genes upregulated in *sod-2* mutants. One of the CEBP-1 target genes that is upregulated in *sod-2* worms is *sqst-1,* which encodes a selective autophagy receptor involved in the degradation of protein aggregates and dysfunctional mitochondria among other targets. SQST-1 has been previously shown to have a role in longevity as overexpression of *sqst-1* increases lifespan in *C. elegans* (39), while overexpression of ref(2)P/dp62, the *Drosophila* homolog of *sqst-1/p62*, increases the lifespan of female flies when initiated at mid-life (40). Also among the CEBP-1 target genes that are upregulated in *sod-2* worms were multiple genes involved in innate immunity including F56D2.5, K09D9.1 and T19D12.4. As enhanced immunity has been associated with extended longevity, the upregulation of these innate immunity genes may also be contributing to lifespan extension in *sod-2* mutants. In addition, CEBP-1 has been shown to bind to the promoter region of transcription factors that mediate different pathways of cellular resilience including the endoplasmic reticulum unfolded protein response (*ire-1*), the p38-mediated innate immune signaling pathway (*sek-1*) and the SKN-1-mediated oxidative stress response (*skn-1*), all of which have been shown to affect longevity (reviewed in (30)).

An important question remains as to how cytoplasmic kinases are activated by increased mitochondrial superoxide when superoxide is not able to cross a biological membrane. One possibility is that superoxide is converted to hydrogen peroxide by either *sod-3* in the mitochondrial matrix, *sod-1* in the intermembrane space or through a non-catalyzed reaction. Since hydrogen peroxide can cross biological membranes, it could transmit the elevated superoxide signal outside of the mitochondria and initiate cytoplasmic signaling cascades through oxidative modification of kinases or other signaling molecules. Another possibility is that the increase in mitochondrial superoxide results in the activation of the mitochondrial unfolded protein response (mitoUPR), or similar pathways that begin in the mitochondria, which then activate cytoplasmic signaling pathways. The mitoUPR has been shown to be activated by ROS (41, 42) and is activated in *sod-2* mutants (**Figure S8**)(43).

Evolutionarily, a programmed genetic response against mitochondrial superoxide may have evolved as a defense mechanism against bacterial pathogens. *C. elegans* generate ROS using an NADPH oxidase/dual oxidase (BLI-3/DUOX) to kill infecting bacteria (44, 45). Bacterial pathogens can cause damage to the mitochondria that results in elevated ROS production (46). In addition, some bacteria can generate ROS during an infection (47–49). Thus, there are multiple mechanisms by which bacterial infections can lead to elevated ROS (11). In order to survive the infection by bacterial pathogens, *C. elegans* may have utilized the SEK-3/PMK-3/MAK-2/CEBP-1 signaling pathway to activate genetic pathways to protect against the infection including genes involved in innate immunity and cuticle formation, which are upregulated in *sod-2* worms, previously associated with longevity and would serve to protect against bacterial pathogens.

## Conclusion

In this work, we find that long-lived *sod-2* mutants exhibit several changes in gene expression including the activation of pathways involved in innate immunity and cuticle development. In a targeted RNAi screen, we identified the kinase MAK-2, as being specifically required for the extended lifespan of *sod-2* worms. We also found that other proteins that act with MAK-2 in established signaling pathways including SEK-3, PMK-3, and CEBP-1 are also specifically required for *sod-2* longevity. Overall, this work provides support for a mitochondria-to-nucleus signaling pathway that promotes longevity in response to elevated mitochondrial superoxide levels.

## Methods

### Strains

The *C. elegans* strains were obtained from the *Caenorhabditis* Genetics Center (CGC) and all strains were outcrossed to N2 a minimum of three times. Strains: N2 – wild-type, MQ1451 *sod-2(ok1030),* JVR649 *mak-1(ok29870),* JVR6652 *sod-2(ok1030);mak-1(ok2987),* JVR650 *mak-2(gk1110),* JVR653 *sod-2(ok1030);mak-2(gk1110),* JVR652 *mnk-1(ok2784),* JVR654 *sod-2(ok1030);mnk-1(ok2784)*.

### RNA isolation and sequencing

To prepare worms for RNA-seq, wild-type and *sod-2* worms were bleached and the eggs were transferred to 10 cm NGM plates seeded with 5X concentrated OP50 bacteria. For the day 8 time point, worms were transferred to NGM plates containing 50 µM FUdR prior to the start of progeny production (late L4 to early young adult stage). Worms were transferred in M9 buffer using a wide pipet tip. When worms reached day 1 or day 8, they were collected in M9 buffer into a 15 ml conical tube. The tube was centrifuged for 2 minutes at 200 g and the supernatant was removed leaving 100-150 µL of buffer. After washing three times with M9 buffer, 1 ml of Trizol was added and the worms were flash frozen in liquid nitrogen. After storage at −80°C, worms were thawed and transferred to Bashing Beads Lysis Tubes (Zymo Research) and vortexed at 3000 RPM. The tubes were then centrifuged at 12,000 g for 1 minute. After lysis, RNA was isolated using the Direct-zol RNA Miniprep kit according to the manufacturer’s instructions. Once the RNA was isolated, RNA sequencing was performed by Genome Quebec. A polyA enriched library was prepared and sequenced on an Illumina NovaSeq X Plus sequencing machine with 25 million reads per sample (PE100). Sequencing metrics can be found in **Table S2**. RNA-seq data has been deposited at NCBI GEO: GSE286219.

### Bioinformatic analysis

Adaptor sequences and low-quality score bases (Phred score < 30) were first trimmed using *Trimmomatic* (50). The resulting reads were aligned to the WBcel235 assembly of the *C. elegans* reference genome, using *STAR* (51). Read counts were obtained using *HTSeq* (52) with parameters *-m intersection-nonempty-stranded=reverse*. For all downstream analyses, we excluded lowly-expressed genes with an average read count lower than 10 across all samples. Batch effects were corrected using ComBat-seq (53). Raw counts were normalized using *edgeR*’s TMM algorithm (54) and were then transformed to log2-counts per million (log2CPM) using the *voom* function implemented in the *limma* R package (55). In comparing gene expression in the three *sod-2* samples from day 1 of adulthood to our previous RNA-seq data from day 1 of adulthood (NCBI GEO: GSE93724), one sample did not cluster with the other 8 samples. This outlier sample was removed and replaced by our six previous *sod-2* day 1 adult samples in identifying differentially expressed genes. To assess differences in gene expression levels, we fitted a linear model using the *lmfit* function. Nominal p-values were corrected for multiple testing using the Benjamini-Hochberg method. Gene set enrichment analysis based on pre-ranked gene list by t-statistic was performed using the R package *fgsea* (http://bioconductor.org/packages/fgsea/). Weighted Venn diagrams were generated using BioVenn (https://www.biovenn.nl/). Gene ontology analyses were performed using GOrilla (https://cbl-gorilla.cs.technion.ac.il/) and ShinyGO 0.81 (https://bioinformatics.sdstate.edu/go/).

### RNA interference lifespan screen

For the RNAi lifespan screen, RNA interference bacteria was grown in 2YT media containing 25 µg/mL Carbenicillin for 16 hours at 37°C on a shaker set at 250 RPM. The bacteria was concentrated 2x and immediately used to seed plates. RNAi plates were freshly made from 35 mm NGM plates containing 1 mM IPTG and 25 µg/mL Carbenicillin. Plates were left to dry for 24 hours before seeding, at which point each plate was seeded with 2x bacteria. Plates were left on benchtop until dry, and then stored at 4°C until needed. Gravid adult worms were bleached and eggs were seeded onto RNAi plates. When worms reached the L4 stage they were transferred to a new RNAi plate. Worms were transferred to new RNAi plates every day until egg laying ceased in order to prevent progeny contamination. Afterwards, worms were transferred on to new RNAi plates as needed. Worms were checked 3 times a week for survival. Survival was initially checked based on observable movement or pharyngeal pumping. If the worm did not show any signs of this, it was gently poked on the tail followed by the head to initiate movement. If the worm still did not move after this, it was marked as dead. Worms that crawled off the plate, died from expulsion of internal organs or internal hatching were censored from the dataset. All worms were maintained and scored at 20°C. Raw lifespan data can be found in **Table S3**.

### Double mutant lifespan

Lifespan assay was conducted at 20°C following a similar protocol to that described above except the regular NGM plates were used instead of RNAi plate. Regular NGM plates were made in 35 mm petri dishes and seeded with 2x OP50 grown in LB for 20 hours.

### Lifespan with 0.1 mM paraquat

NGM plates were prepared with 0.1 mM paraquat. 50 µM FUdR was also included in the plates to inhibit the development of progeny and to prevent internal hatching of progeny (56). Plates were seeded with 10X concentrated OP50 bacteria as paraquat inhibits the growth of OP50. To start the assay, pre-fertile young adult worms were transferred to either 0.1 mM paraquat, 50 µM FUdR plates or control plates containing just 50 µM FUdR. Worms were transferred to fresh plates after 4-5 days and as needed thereafter. Survival of the worms was checked every 2-3 days until all of the worms were dead. A minimum of three biological replicates were completed per strain.

### Brood size

Brood size was determined by placing individual L4 worms on NGM plates. Worms were transferred daily to new plates until egg laying ceased. The progeny was allowed to develop up to at least the L3 stage before quantification. Three biological replicates were performed for each strain, with each biological replicate containing at least four worms.

### Development time

Developmental time was assessed by allowing gravid adult worms to lay eggs. Three biological replicates were performed for each strain. For strains N2, *mak-1, mak-2* and *mnk-1*, 30 gravid worms for each were allowed to lay eggs for 30 minutes. For *sod-2,* 30 worms were allowed to lay eggs for 4 hours, while the strains *sod-2;mak-1, sod-2;mak-2* and *sod-2;mnk-1* had 90 worms allowed to lay eggs for 2 hours. After the allotted egg laying time, the gravid worms were removed. The hours from egg laying to adult was measured as the developmental time.

### Thrashing rate

Young adult worms were transferred to 60 mm NGM plates seeded with OP50 *E. coli*. A volume of 1 mL of M9 buffer was added to the worms. The worms were left to acclimatize for 1 min before capturing 1 min video at 14 FPS using WormLab (MBF Bioscience). The videos were analyzed using wrMTrck plugin for the open-source image processing software, Fiji. Worm tracks were only considered for analysis when they included at least 420 frames (30 seconds). The results were pooled from three biological replicates.

### Heat stress resistance

OP50 bacteria was grown in LB at 37°C overnight and was then concentrated 10x. Freshly made 60 mm NGM plates were seeded with 75 µL of 10X concentrated OP50 bacteria. Seeded plates were allowed to dry at 35°C for ∼15 minutes. Once dried, young adult worms were transferred to the plates and incubated at 35°C for 6 hours, after which plates with worms were transferred to 20°C. Scoring for survival occurred 24 hours after the start of the heat shock (24 hours after the worms were placed into the 35°C). Three biological replicates were completed for each strain.

### Osmotic stress resistance

NGM plates containing 450 mM NaCl were poured and allowed to dry on the bench before storing at 4°C. One day before the experiment, 200 µL of 5X concentrated OP50 *E. coli* was seeded onto each plate and allowed to dry overnight. The next day, young adult worms were picked into the NGM plates containing 450 mM NaCl and incubated at 20°C for 48 h before scoring survival. Worms with internal hatching were censored and excluded from the total number of deaths. Three biological replicates were completed for each strain.

### Resistance to acute oxidative stress

Resistance to acute oxidative stress was assessed by exposing worms to NGM plates containing 320 µM juglone. Juglone was freshly dissolved in ethanol. Freshly made NGM solution was made, autoclaved and allowed to cool to below 60°C at which point the juglone was aliquoted into the NGM solution. Afterwards, the solution was poured into 60 mm plates and allowed to dry for 10-15 minutes. Once dry, 20 µL of 10X OP50 was pipetted on to each plate and spread across the plate using a Pasteur glass pipette with a bent tip. The plates were allowed to dry for another 10 minutes, at which point young adult worms were transferred to each pate. The worms were scored for death every 2 hours up to the 8-hour mark. A total of three biological replicates were performed for each strain.

### Resistance to chronic oxidative stress

Resistance to chronic oxidative stress was assessed by exposing worms to 4 mM paraquat (Methyl viologen dichloride hydrate). Paraquat freshly dissolved in ddH_2_O was added to NGM before pouring and 100 µM FUdR was included in the plates to prevent internal hatching that is caused by exposure to paraquat. The plates were allowed to dry overnight before storage at 4°C. One day before the experiment, the paraquat plates were seeded with 200 µL of 10X concentrated OP50 *E. coli* and allowed to dry overnight. The next day, young adult worms were transferred to these plates and survival was monitored every day until death. Three biological replicates were completed.

### Anoxia resistance

Freshly made 60 mm NGM plates containing 50 µM FUdR were seeded with 250 µL of 2x concentrated OP50 and let dry overnight on bench top. Once dry, plates were stored at 4°C until needed. Young adult worms from each strain were placed on to the plates, afterwards the plates were placed into BD GasPak™ EZ Anaerobe (Cat. 260683) and incubated for 48 hours at 20°C. Subsequently, the bags were unsealed and allowed to reoxygenate for 24 hours at 20°C, at which point the worms were scored for survival. Three biological replicates were completed for each strain.

### Bacterial pathogen resistance

Resistance to bacterial pathogen stress was assessed using *P. aeruginosa* strain PA14. The slow kill assay was performed as previously described (9, 57). Briefly, young adult worms were grown on plates containing 2x concentrated OP50 and 100 mg/L FUdR until day 3 of adulthood. An overnight PA14 culture was seeded onto plates containing 20 mg/L FUdR and the bacteria was allowed to grow for one day at 37°C followed by one day at 20°C. The Day 3 adult worms were then transferred to the PA14 plates and survival was monitored daily. Three biological replicates were performed with approximately 40 worms per replicate.

### Construction of RNAi clones

As the RNAi clone of *mkk-4* (WBGene00003368) was not available from Ahringer *C. elegans* RNAi feeding library or *C. elegans* ORFeome (Vidal) Library, a new RNAi feeding clone was generated in our laboratory. Through nested PCR, a cDNA library from Day 1 adults was used to amplify the cDNA of *mkk-4* using Platinum SuperFi II Green PCR Master Mix with the following primers (TCA ACA CGT CCC ACA TCA CT and CTC CAA GAA TTC GCT TAG CC). The product PCR was further diluted 1:100 in ddH_2_O and was amplified again using the following primers (GGT GGT GGT ACC AAA CGG CAA TTT TGG AAC AG and GGT GGT ACT AGT TCG TCG CTG TCT GGA TGT AG) which flank the ORF with KpnI and SpeI cutting sites resulting in a 693 bp fragment. The product PCR was purified using EZ-10 Spin Column PCR Products Purification Kit. The purified PCR and the empty L4440 plasmid were double digested using KpnI-HF and SpeI-HF (New England Biolabs) and ligated using Instant Sticky-end Ligase Master Mix (New England Biolabs) according to the manufacturer instructions. The ligated product was transformed into chemically competent HT115(DE3) *E. coli* and the cells were spread on LB plate containing 100 µg/mL carbenicillin and 10 µg/mL tetracycline and allowed to grow overnight at 37°C. The resultant colonies were sequenced verified using M13F primer to ensure correct insert.

### Statistical analysis

Statistical analysis was performed using GraphPad Prism version 9. Statistical tests utilized are indicated in the figure legends. Error bars indicate standard error of the mean (SEM). ns=not significant, *p<0.05, **p<0.01, ***p<0.001, ***p<0.0001.

## Acknowledgments

Some strains were provided by the CGC, which is funded by NIH Office of Research Infrastructure Programs (P30 OD010440). We would also like to acknowledge the *C. elegans* knockout consortium and the National Bioresource Project of Japan for providing strains used in this research. This work was supported by the Canadian Institutes of Health Research (CIHR; http://www.cihr-irsc.gc.ca/; JVR; Application 399148 and 416150) and the Natural Sciences and Engineering Research Council of Canada (NSERC; https://www.nserc-crsng.gc.ca/index_eng.asp; JVR; Application RGPIN-2019-04302). JVR is the recipient of a Senior Research Scholar career award from the Fonds de Recherche du Québec Santé (FRQS) and Parkinson Quebec. The funders had no role in study design, data collection and analysis, decision to publish, or preparation of the manuscript.

## Author Contributions

Conceptualization: UA, JVR. Methodology: UA, ATG, MMP, AA, JVR. Investigation: UA, ATG, MMP, AA. Analysis: UA, ATG, MMP, AA, AP, JVR. Validation: UA, ATG, MMP, AA, JVR. Visualization: UA, ATG, MMP, AA, AP, JVR. Writing – original draft: UA, JVR. Writing – review and editing: UA, ATG, MMP, AA, AP, JVR. Supervision: JVR.

## Data and materials availability

Raw data will be provided upon request. All materials used in this manuscript are available to be shared with the scientific community. Requests for data or materials should be addressed to Jeremy Van Raamsdonk.

## Supplemental Figures

**Figure S1.**
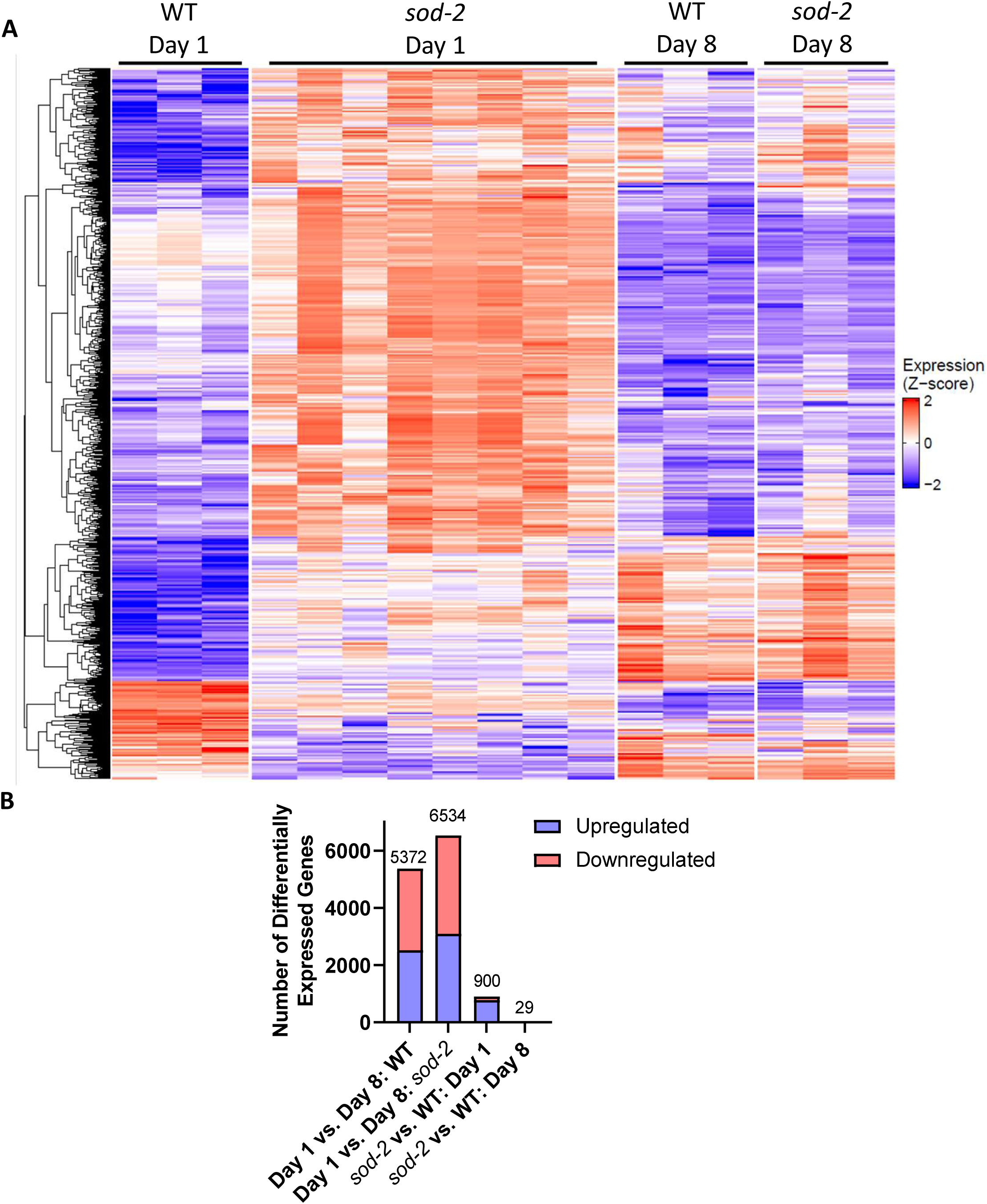
Differences in gene expression between time points are much greater than differences between *sod-2* and wild-type. **(A)** A heat map shows marked changes in gene expression between *sod-2* and wild-type worms at day 1 of adulthood and between day 1 and day 8 worms. Gene expression between the genotypes was more similar at day 8 of adulthood. **(B)** The number of differentially expressed genes between day 1 and day 8 of adulthood was much greater than the number of differentially expressed genes between *sod-2* and wild-type worms.

**Figure S2.**
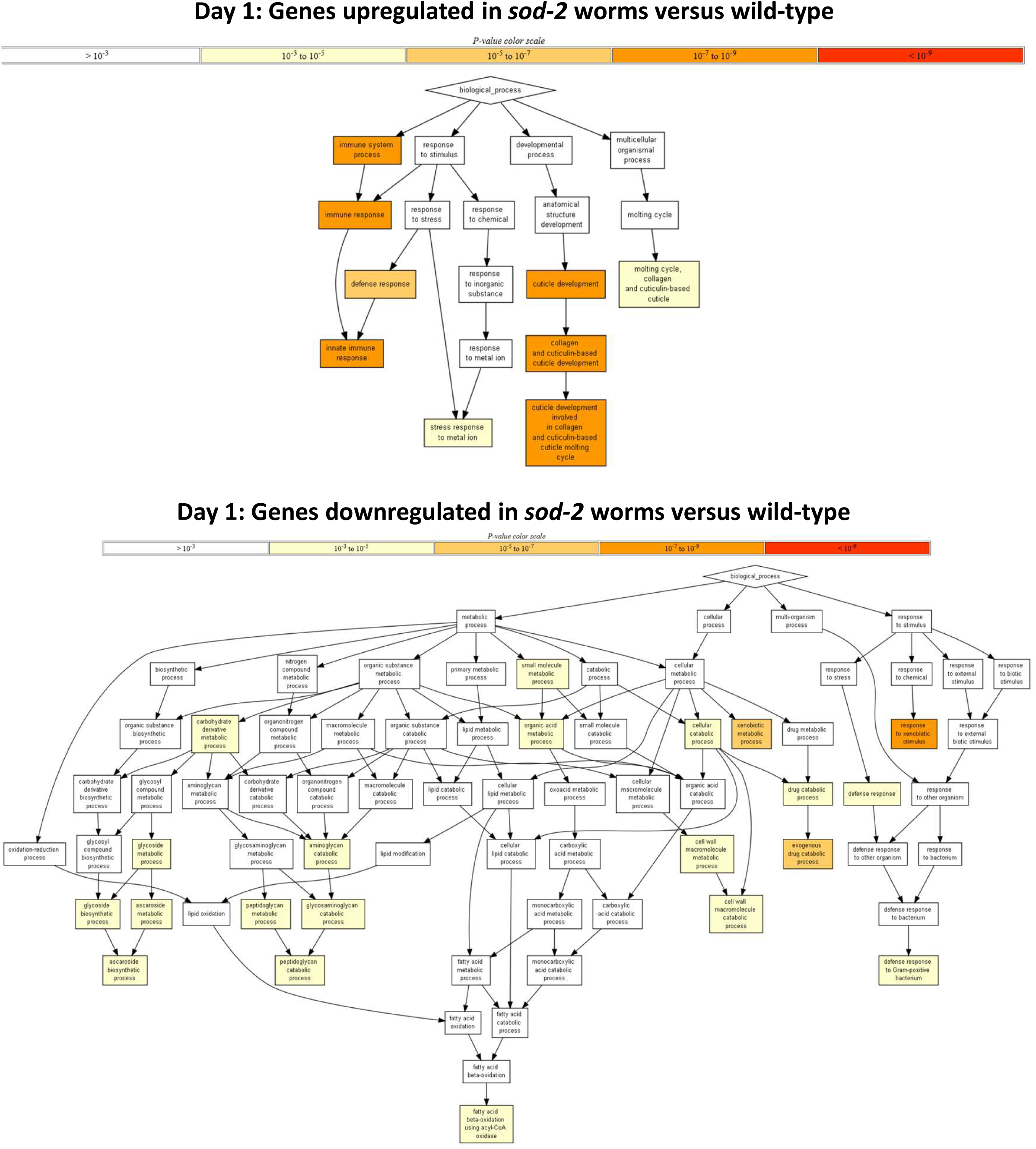
Enrichment analysis of differentially expressed genes in *sod-2* worms. Differentially expressed genes were analyzed to identify overrepresented gene ontology (GO) groups using Gene Ontology enRIchment anaLysis and visuaLizAtion tool (GOrilla: https://cbl-gorilla.cs.technion.ac.il/). At day 1 of adulthood, *sod-2* worms exhibit a significant upregulation of genes involved in cuticle structure and innate immunity.

**Figure S3.**
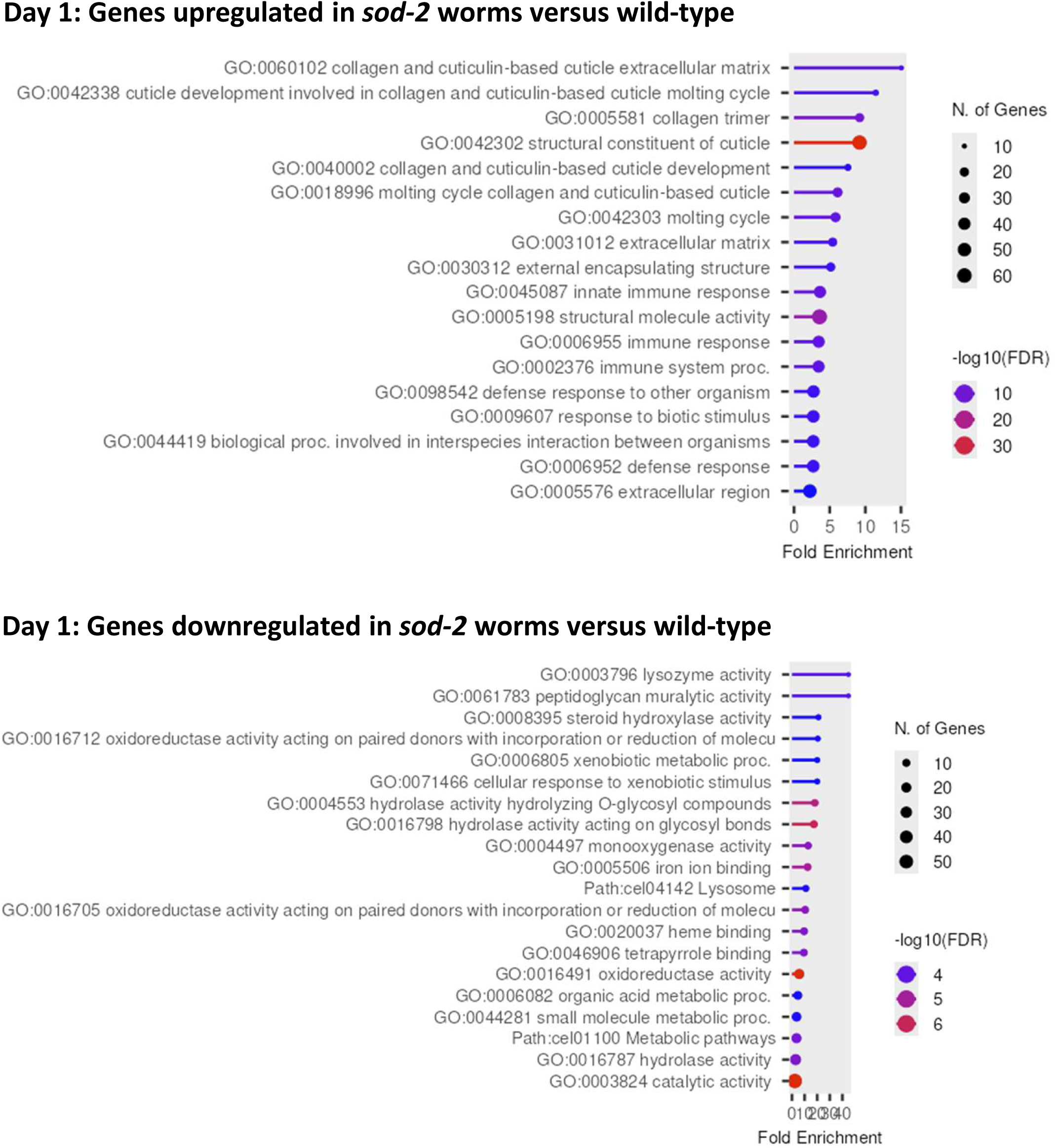
Enrichment analysis of differentially expressed genes in *sod-2* worms. Differentially expressed genes were analyzed to identify overrepresented gene ontology (GO) groups using ShinyGO 0.81 (https://bioinformatics.sdstate.edu/go/). At day 1 of adulthood, *sod-2* worms exhibit a significant upregulation of genes involved in cuticle structure and innate immunity.

**Figure S4.**
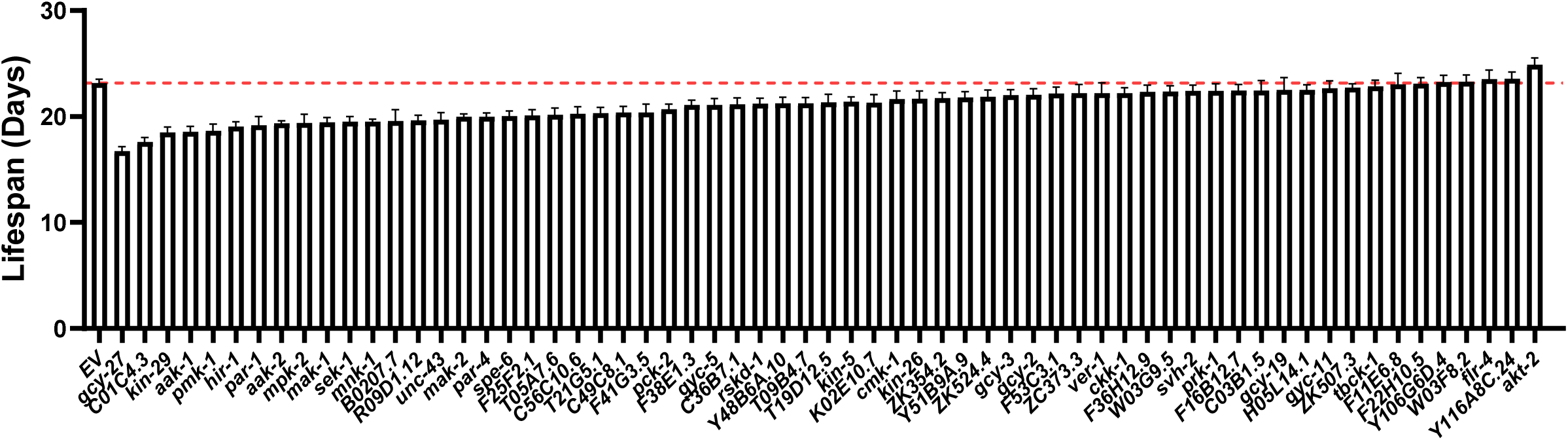
Full results of kinase-targeted RNAi lifespan screen. All lifespans were performed in *sod-2* mutants. Error bars indicate SEM. A minimum of three biological replicates were performed. Red line indicates lifespan on empty vector (EV). The results from RNAi clones that were significantly different from EV are shown in Figure 2C.

**Figure S5.**
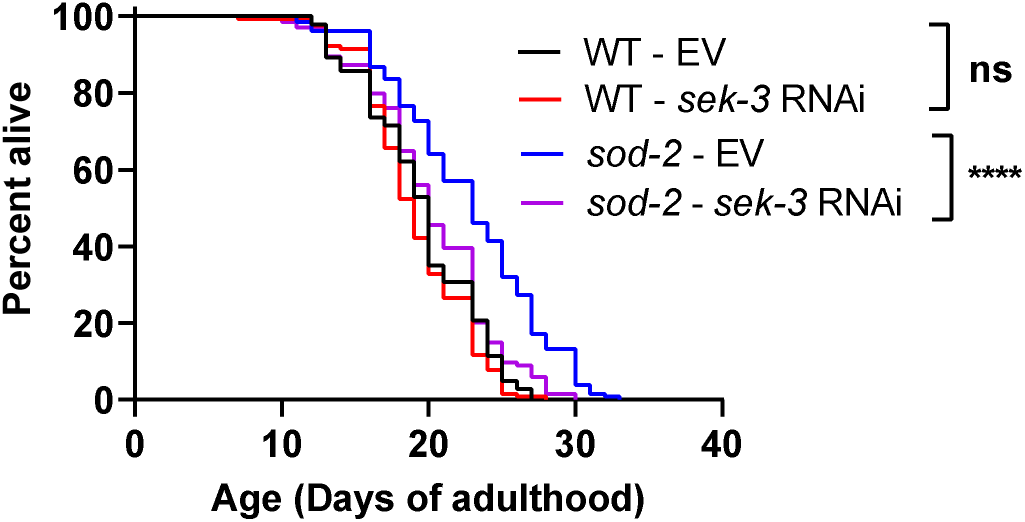
*sek-3* is required for lifespan extension in *sod-2* worms. Knocking down *sek-3* using RNAi decreased *sod-2* lifespan but had no effect on wild-type longevity. Three biological replicates were performed. Statistical significance was assessed using a log-rank test.

**Figure S6.**
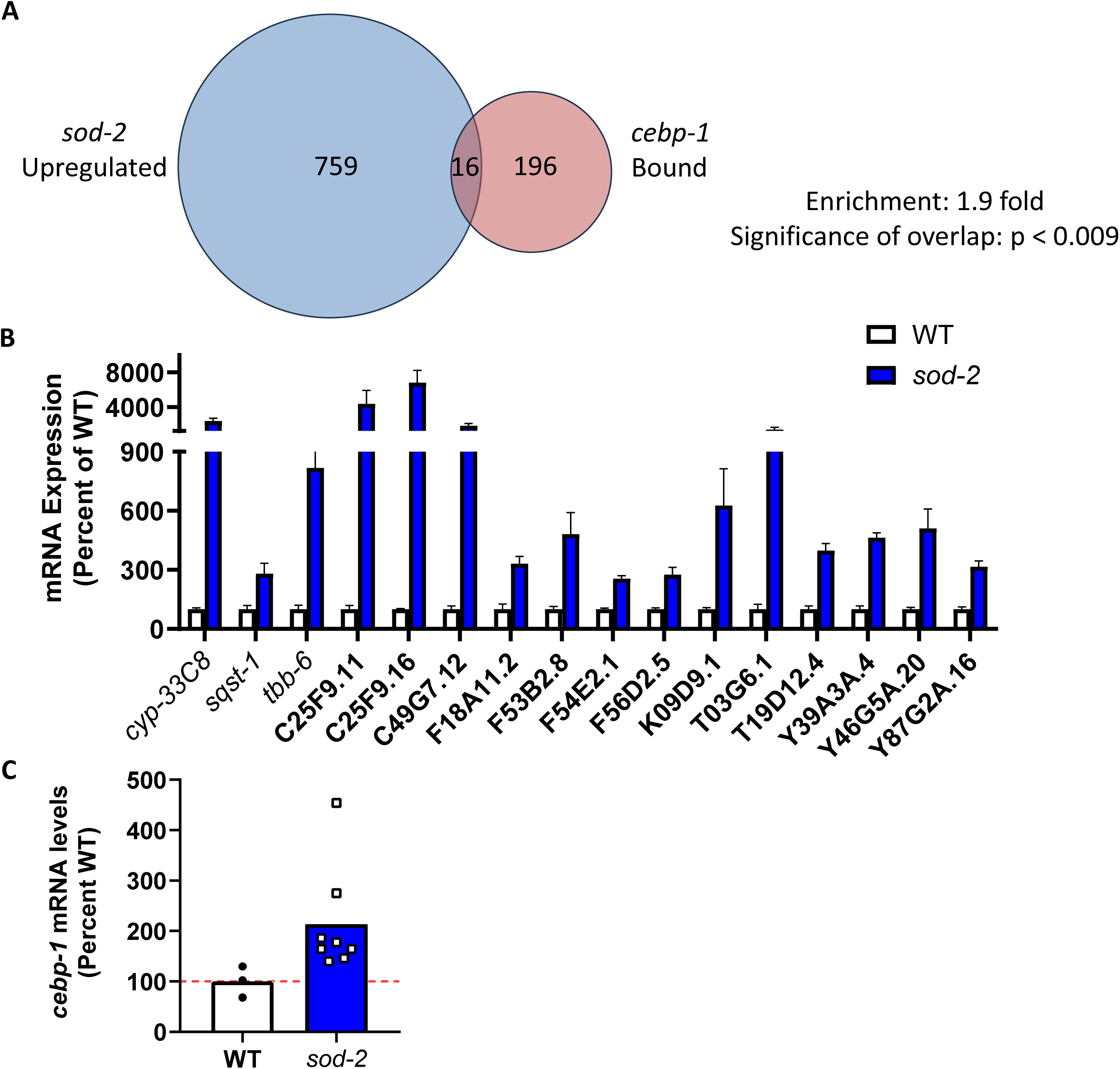
Enrichment of CEBP-1 target genes among genes significantly upregulated in *sod-2* worms. **(A)** Genes upregulated at day 1 of adulthood in *sod-2* worms compared to wild-type worms were compared to genes bound by CEBP-1 in a previously published ChIP-seq experiment. There were 16 genes in common, which represents a statistically significant enrichment compared to the number expected if all of the genes were picked at random. **(B)** Bar chart showing expression levels of the 16 overlapping genes in *sod-2* worms versus wild-type worms at day 1 of adulthood. **(C)** *sod-2* worms exhibit a trend towards increased expression of *cebp-1.* Error bars indicate SEM.

**Figure S7.**
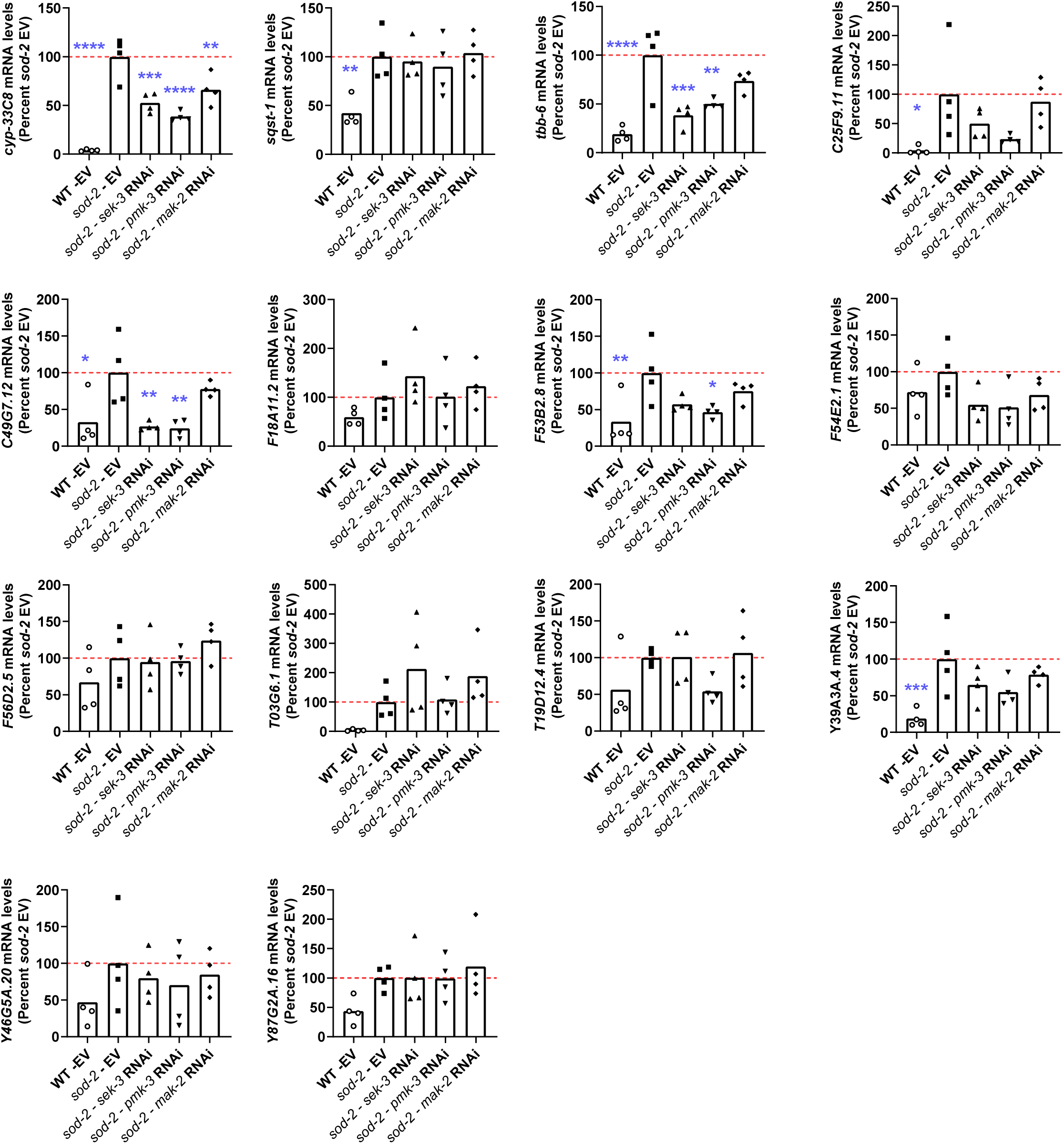
SEK-3/PMK-3/MAK-2 pathways contributes to upregulation of CEBP-1 target genes in *sod-2* mutants. Worms were treated with RNAi targeting the kinases *sek-3, pmk-3* or *mak-2.* The expression of CEBP-1 target genes was measured using quantitative RT-PCR. Expression levels were compared to empty vector (EV) treated control worms. For all of the genes tested, we observed increased expression of the CEBP-1 target gene in *sod-2* mutants on EV compared to wild-type worms on EV. The expression of multiple CEBP-1 target genes was significantly reduced by knocking down *sek-3, pmk-3* or *mak-2.* This indicates that the SEK-3/PMK-3/MAK-2 pathway contributes to the upregulation of these genes in *sod-2* mutants. Three biological replicates were performed. Statistical significance was assessed using a one-way ANOVA with Dunnett’s multiple comparisons test. Statistically significant differences from the *sod-2 –* EV group are indicated.

**Figure S8.**
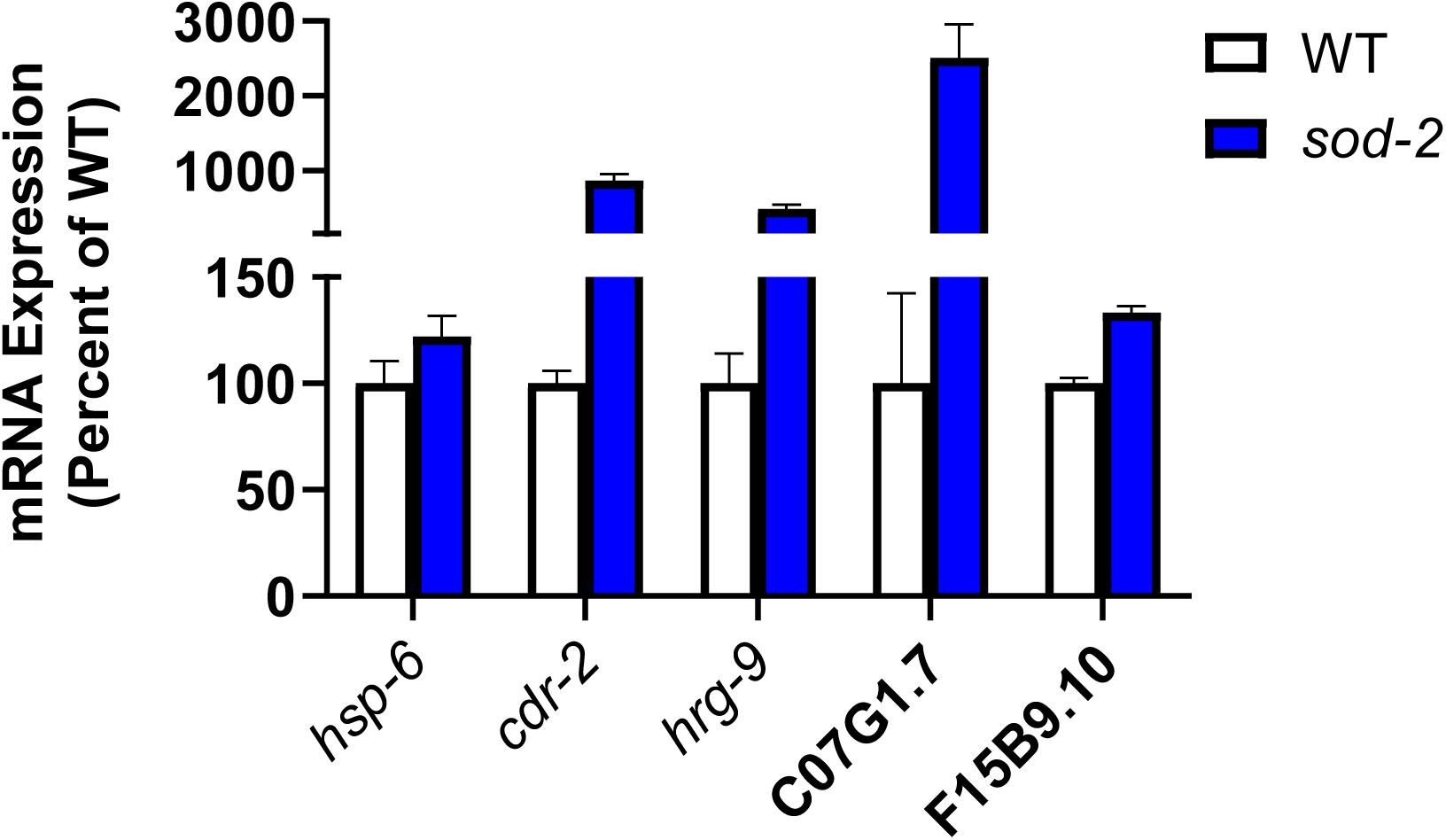
Activation of the mitochondrial unfolded protein response in *sod-2* mutants. The expression of high confidence mitoUPR target genes was examined in *sod-2* mutants from RNA-seq data on day 1 of adulthood. All of the mitoUPR target genes showed increased levels in *sod-2* mutants compared to wild-type.

